# HTS-Oracle: Experimentally validated AI-enabled prioritization for generalizable small molecule hit discovery

**DOI:** 10.1101/2025.11.26.690784

**Authors:** Hossam Nada, Laura Calvo-Barreiro, Natalie Fuchs, Baljit Kaur, Sungwoo Cho, Saurabh Upadhyay, Nicholas A Meanwell, Moustafa T. Gabr

## Abstract

High-throughput screening (HTS) remains a central pillar of small molecule discovery yet routinely fails for immune receptors and protein-protein interaction-driven targets. Here, we introduce HTS-Oracle, an experimentally validated AI system for prospective hit discovery that integrates molecular language modeling with cheminformatics to prioritize bioactive compounds at scale. We deploy HTS-Oracle across three clinically validated yet historically intractable immune targets, TREM2, CHI3L1, and CD28, representing cryptic binding pockets, intrinsically disordered proteins, and protein-protein interaction-driven immune checkpoint, respectively. Across the tested targets, HTS-Oracle reduces experimental screening requirements by up to >99% while increasing hit rates by up to 176-fold relative to traditional HTS. Notably, the platform remains predictive under extreme data sparsity, achieving an eightfold improvement for CD28 despite fewer than 2% actives in training. By consistently enriching for experimentally validated hits, HTS-Oracle establishes a new performance benchmark for hit discovery and unlocks small molecule access to immune targets long regarded as chemically inaccessible.

---

High-throughput screening (HTS) remains a cornerstone of early-phase small molecule drug discovery, serving as the primary strategy for identifying initial chemical hits across therapeutic areas^1–3^. An analysis of 58 approved drugs showed that approximately 30% originated from HTS campaigns, and this approach remains the predominant method for hit identification in many leading pharmaceutical companies^4^. Over the years, HTS has undergone remarkable technological evolution, with innovative platforms such as temperature-related intensity change (TRIC)^5^, surface plasmon resonance (SPR)^6^, and mass spectrometry-based approaches^7^ enhancing the speed, resolution, and diversity of hit detection.

Despite technological advancements, conventional HTS campaigns often suffer from high costs, limited efficiency, and low success rates—typically yielding hit rates of just 1–2%^8,9^. These inefficiencies are especially pronounced when screening against targets with limited tractability to small molecules, such as immune receptors and protein–protein interaction (PPI) interfaces. In addition to high attrition, HTS workflows often discard negative screening data, eliminating valuable information that could inform future predictions and compound prioritization^10,11^. As a result, vast amounts of experimental data remain underutilized, and chemical space exploration remains largely unguided.

Artificial intelligence (AI) and machine learning (ML) offer a powerful opportunity to address these challenges by enabling large-scale virtual screening and compound prioritization^12^. Deep learning models have shown success in capturing nonlinear structure–activity relationships and accelerating hit discovery across vast chemical spaces^13,14^. However, many ML frameworks are designed for retrospective performance benchmarking, lack integration with real-world HTS pipelines, and fail to incorporate negative data, limiting their prospective utility in challenging screening contexts. To overcome these limitations, we developed HTS-Oracle, a retrainable, multi-modal deep learning platform that integrates transformer-derived molecular embeddings with traditional cheminformatics descriptors. HTS-Oracle combines ChemBERTa^15^, a SMILES-based language model, with RDKit-derived molecular fingerprints and physicochemical features in an ensemble architecture optimized for prospective hit prediction. By uniting contextual and structural representations through multiple feature selection strategies, HTS-Oracle prioritizes high-confidence hits from large compound libraries. Unlike traditional traditional quantitative structure–activity relationship (QSAR) models, HTS-Oracle is explicitly designed for prospective screening, retraining, and deployment across new targets, thereby enabling rapid adaptation and reuse. HTS-Oracle was trained using ChEMBL (public repository) and scientifically published data, followed by an evaluation for three distinct and key therapeutic targets: Triggering Receptor Expressed on Myeloid Cells 2 (TREM2), Chitinase-3-like protein 1 (CHI3L1) and Cluster of Differentiation 28 (CD28). These immune-related proteins are implicated in Alzheimer’s disease, cancer, and autoimmune disorders, yet have proven highly challenging for conventional small molecule discovery approaches and falls into the category of “undruggable” targets^22–27^. The selection of TREM2, CHI3L1, and CD28 as case studies was driven by several key considerations. First, these proteins represent clinically validated therapeutic targets implicated in major diseases yet remain intractable to traditional small molecule drug discovery. By focusing on these challenging targets, we aimed to demonstrate the ability of HTS-Oracle to overcome key limitations of conventional HTS. In addition, the three chosen targets have been highly implicated in various disorders making them highly relevant for clinical development. The successful identification of potent leads for TREM2 and CHI3L1, including compounds with improved binding affinities compared to known ligands, demonstrates the capability of HTS-Oracle in accelerating drug discovery for targets with limited small molecule tractability.

Additionally, these case studies were strategically chosen to evaluate HTS-Oracle’s performance under varying data constraints which are typical of early-stage screening campaigns particularly regarding dataset balance and origin. Training datasets for the three target-specific models were compiled from public chemical databases and published high-throughput screening studies. These datasets exhibited substantially different degrees of class balance, with hit rates ranging from severe imbalance in CD28 (<2% active compounds) to moderate balance in TREM2 (14% active compounds) and relative balance in CHI3L1 (48% active compounds). The performance of HTS for all three targets exceeded the expectations and was significantly higher than traditional HTS campaigns for the same targets.

Comparative analysis across the three therapeutic targets (Figure 1) demonstrates that HTS-Oracle substantially reduces experimental burden while improving hit rates. The platform decreased the number of compounds requiring experimental validation by >99% for TREM2 and CHI3L1 and by 70% for CD28, highlighting its capacity to prioritize hits with minimal wet-lab effort. Correspondingly, enrichment factors ranged from 8.4× to 176× relative to traditional HTS, underscoring robust target-specific performance even under challenging data conditions such as the highly imbalanced CD28 training set. These findings indicate that HTS-Oracle provides a scalable, generalizable, and experimentally efficient approach to hit identification, enabling substantial reductions in cost and time while maintaining or improving discovery outcomes compared to conventional screening workflows.

**Figure 1.**
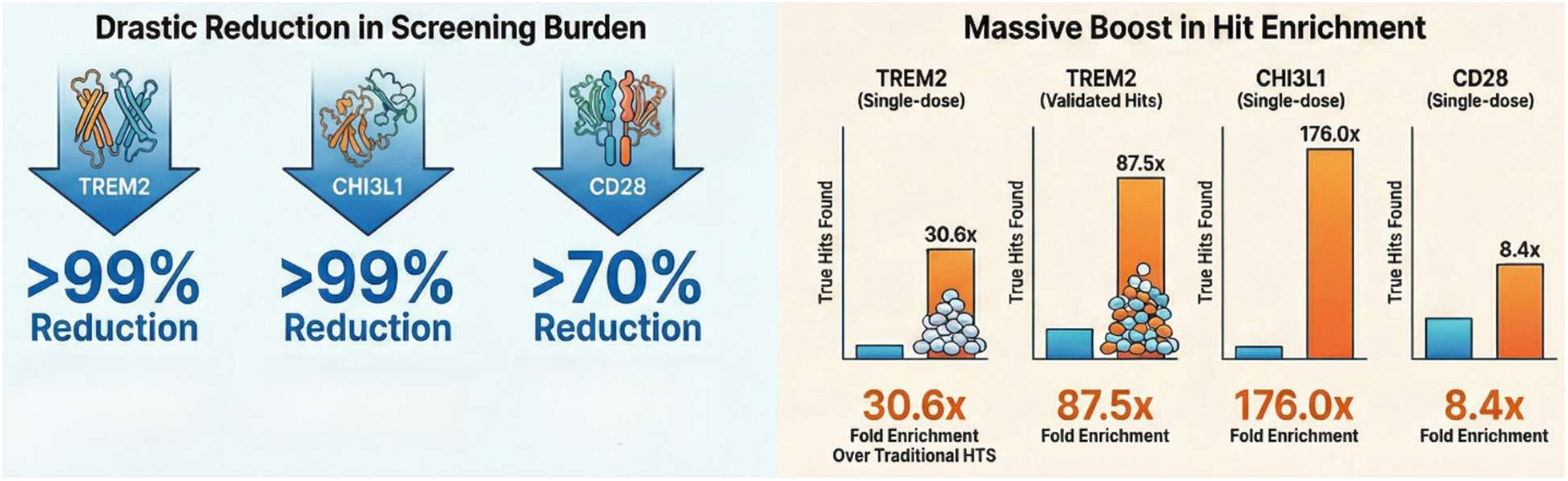
Performance comparison of HTS-Oracle with traditional HTS. **(A) Screening burden reduction:** For three independent targets (TREM2, CHI3L1, and CD28), HTS-Oracle reduced the number of compounds requiring experimental testing by >99% for TREM2 and CHI3L1, and by 70% for CD28, relative to traditional HTS workflows. **(B) Hit rate improvement:** HTS-Oracle achieved substantial increase in hit rate when compared with traditional HTS, including 30.6× and 87.5× for TREM2 (single-dose and validated datasets, respectively), 176× for CHI3L1 (single-dose), and 8.4× for CD28.

## Results and discussion

### Model Architecture, Training and Web Application

A hybrid deep learning model was constructed with a multi-modal ensemble architecture, designed to integrate chemical language modeling with traditional molecular descriptors for binary classification of chemical structures. At the core of this architecture (Figure 2) is a dual-branch neural network that processes two complementary types of molecular representations in parallel. The first branch leverages ChemBERTa^15^, a RoBERTa-based transformer model pre-trained specifically on chemical SMILES strings, to extract contextualized molecular embeddings. The ChemBERTa branch generates 768-dimensional embeddings from the classification [CLS] token, which are regularized using a dropout layer before being input to a multi-layer perceptron (MLP) composed of two linear layers (768→256→128) with batch normalization, ReLU activation, and 30% dropout between layers.

**Figure 2:**
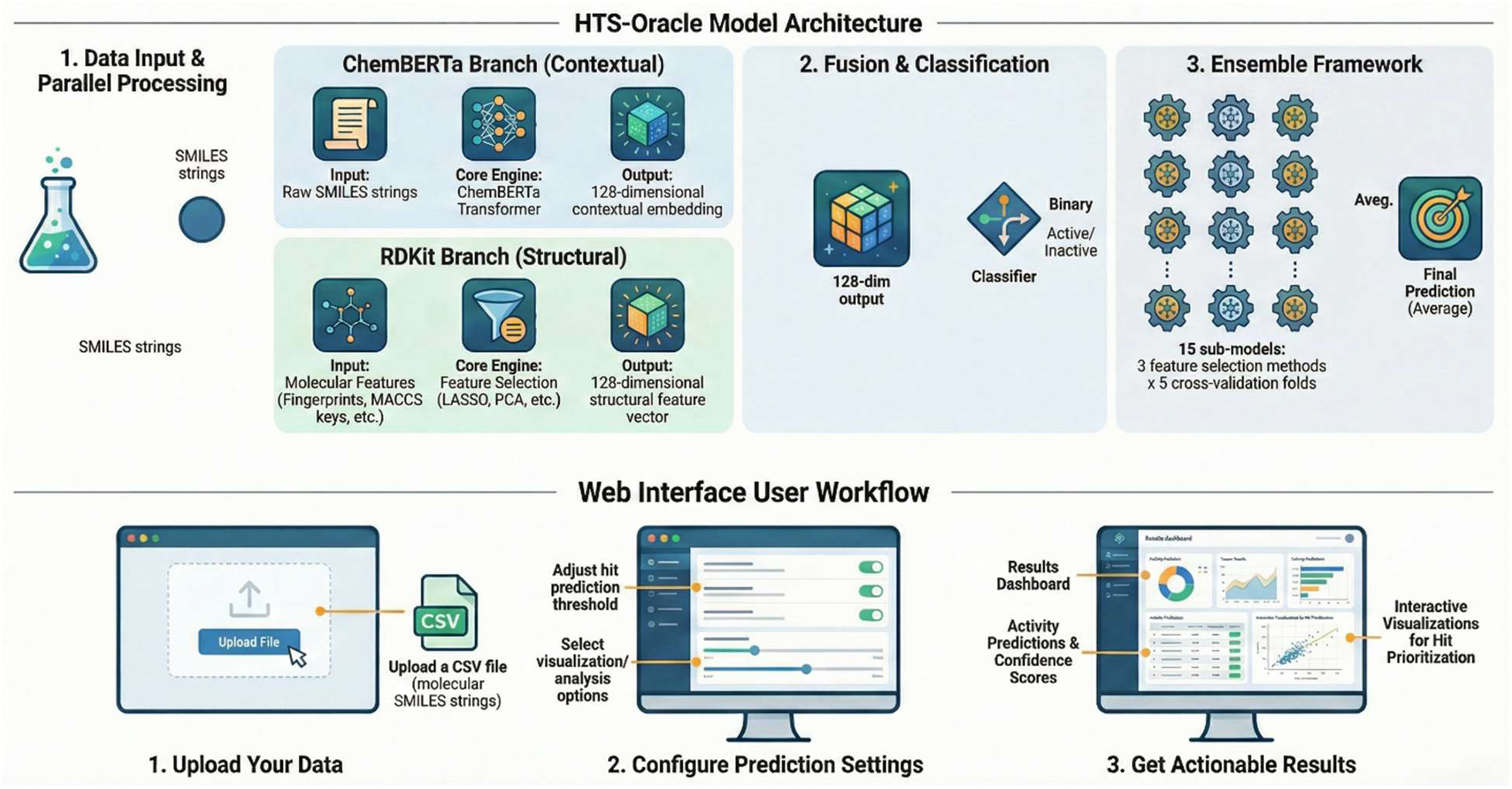
HTS-Oracle architecture (A) and GUI (B). The model features two parallel processing branches that merge *via* late fusion. The HTS-Oracle graphical user interface enables users to upload compound libraries in CSV format, select models and prediction thresholds, and access visualization and reporting options. The interface is designed to provide a streamlined, user-friendly, one-click workflow for hit prediction.

The second branch handles cheminformatics features derived from RDKit^16^ which included 2048-bit Morgan fingerprints, 167-bit MACCS keys, and 15 physicochemical descriptors (e.g., molecular weight, Log*P*, rotatable bonds, hydrogen-bond donors and acceptors, topological polar surface area, ring systems, and drug-likeness metrics). These features undergo dimensionality reduction using one of three methods—LASSO-based feature selection, principal component analysis (PCA), or mutual information filtering which typically reduces the feature space to 200 components. This reduced representation is passed through a parallel MLP with an architecture identical to that of the ChemBERTa branch, allowing harmonized processing of diverse input modalities.

The 128-dimensional outputs from both branches were concatenated into a unified 256-dimensional representation and passed through a final classification head comprising three fully connected layers (256→128→64→1). This classifier used batch normalization and ReLU activations between layers and applied graduated dropout rates (30% and 15%) to encourage generalization and mitigate overfitting. The model employed an ensemble framework by training distinct models for each feature selection method (LASSO, PCA, mutual information) across five stratified folds, yielding a total of 15 sub-models. During inference, predictions from all sub-models were averaged to produce the final ensemble output, enhancing robustness and minimizing overfitting.

To prevent overfitting and minimize bias, the model incorporated multiple regularization techniques, including dropout at multiple stages, batch normalization to reduce internal covariate shift^17^, L2 weight decay *via* the AdamW optimizer, and gradient clipping to stabilize training^18^. Early stopping was employed with a five-epoch patience window based on validation AUC to avoid overtraining. To address the class imbalance inherent in HTS datasets, where hits are sparse, positive class weights were dynamically balanced^19^ for the binary cross-entropy loss based on the class distribution in each training fold. This approach prevented the model from favoring the majority class.

The training pipeline also included several mechanisms to ensure data integrity and model robustness. SMILES strings were validated and parsed with error-handling fallbacks, while molecular features were standardized using StandardScaler to normalize scales across descriptors. NaN and infinite values were automatically replaced with zeros, and safe fallback strategies were used during metric computation to handle edge cases such as single-class predictions. Learning rate scheduling was handled *via* a ReduceLROnPlateau strategy^20^, which adapted the learning rate based on validation performance to avoid premature convergence or overshooting. Finally, the integrity of the validation step was maintained through stratified train–validation splits^21^, ensuring balanced class distributions without leakage. Each fold had an independent validation set to provide unbiased performance estimates.

To simplify the use of the trained models, a web application, named HTS-Oracle, was developed to provide an interactive and scalable platform for predicting molecular activity across compound libraries using the pre-trained ensemble model. At its core, the application integrates multiple feature selection methods (LASSO, PCA, and mutual information) to generate calibrated activity scores for each compound, offering users binary classifications, various visualization options as well as confidence levels. In scenarios where no known actives are available to train a predictive model, HTS-Oracle includes a fallback system that filters libraries based on classical drug-likeness rules. This system applies Lipinski’s rule of five and quantitative estimate of drug-likeness (QED) to rank the compounds based on their predicted absorption, distribution, and overall bioavailability profiles. Collectively, these features deliver an intuitive web-based interface that enables researchers to leverage HTS-Oracle’s predictive capabilities without requiring computational expertise.

Beyond predictions, the application offers a suite of interactive visualizations powered by Plotly and Matplotlib, enabling users to explore the distribution of prediction scores, compare feature selection methods, and correlate predicted activity with key molecular properties. With built-in error handling for invalid SMILES, support for model interpretability, and configurable thresholding, the app serves as a practical tool for virtual screening, hit prioritization, and rational library design in early-stage drug discovery.

### Applications of HTS-Oracle in Small Molecule Drug Discovery

#### Case study 1: HTS-Oracle Achieves 87.5-Fold Enrichment in TREM2 Hit Discovery

The first application of HTS-Oracle involved its training on published TREM2 HTS assay data^28^ and using the trained model to screen an in-house compound library followed by experimental validation of the predicted hits. During training, HTS-Oracle showed consistent performance across all five cross-validation folds, with training loss curves showing smooth convergence by epoch 10 (Figure S1). The validation AUC trajectories reveal some fold-to-fold variability, with Fold 3 achieving the highest peak performance (AUC ≈ 0.88 at epoch 4), while Fold 2 exhibited the most modest performance (AUC ≈ 0.73 at epoch 4) with the overall performance of AUC = 0.83. Comparison of the three feature selection methods within HTS-Oracle (Figure S1) demonstrates distinct prediction behaviors for TREM2 hits. Most notably, the LASSO method exhibited a bimodal distribution with distinct peaks near 0 and 0.8, suggesting identification of patterns related to key pharmacophoric elements essential for TREM2 interaction.

The trained TREM2 HTS-Oracle model was employed to screen an inhouse library comprising 12712 compounds which were purchased from Enamine. The screening resulted in substantial improvements in screening efficiency and hit identification compared to traditional high-throughput screening approaches. In our previous TREM2 screening campaigns using conventional methods with the Dianthus instrument, we achieved hit rates of 1.44% across two focused libraries (TargetMol Lipid Metabolism library, 653 compounds; TargetMol Saccharide and Glycoside library, 595 compounds)^28^. In contrast, HTS-Oracle achieved a remarkable 44% hit rate at the single-dose screening stage (11 hits out of 25 compounds tested). HTS-Oracle required the testing of only 25 compounds versus the >2,500 compounds screened in traditional campaigns which reduced both material costs and experimental time by over 99%. This efficiency gain translates to substantial savings in protein reagents, labeling kits, and consumables, while reducing plate reader occupancy from multiple hours to approximately 10 minutes—a critical consideration for high-demand shared instrumentation.

A key challenge in traditional HTS workflows is the high false-positive rate observed in single-dose screening assays^28–30^ which we previously documented in our TREM2 campaigns. Our conventional approach showed a 4.5-fold attrition rate when transitioning from single-dose screening (1.44% hit rate) to dose-response validation studies (0.32% confirmed hit rate)^28^ primarily due to non-specific binding artifacts and other assay interferences at single concentrations. To evaluate whether HTS-Oracle predictions would exhibit similar attrition, we subjected all 11 initial hits to rigorous dose-response binding affinity measurements using Monolith X across a broad concentration range (5 nM to 200 µM). Notably, seven of the 11 predicted hits (63.6%) demonstrated dose-dependent TREM2 binding with well-defined binding isotherms, yielding a validated hit rate of 28% (7/25 compounds tested). This represents an 87.5-fold enrichment compared to the 0.32% validated hit rate from our previous traditional screening campaigns. The superior performance of HTS-Oracle predictions in dose-response studies (Figure 3 and Figure S2) suggests the platform effectively learns structural features associated with specific, high-quality target engagement rather than promiscuous binding behavior. Furthermore, the validated HTS-Oracle hits exhibited improved binding affinities compared to compounds identified through conventional screening.

**Figure 3.**
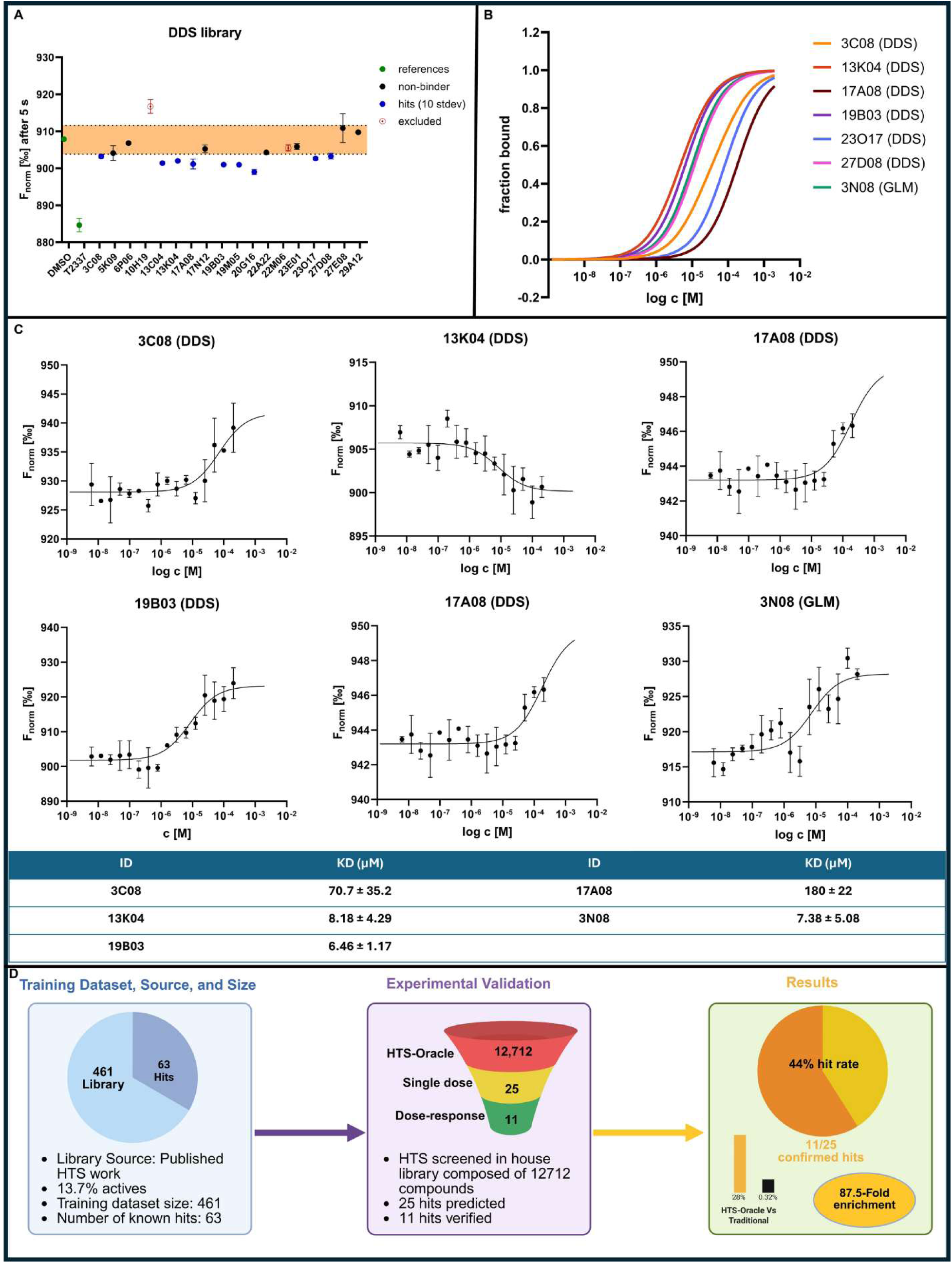
Experimental validation and binding characterization of HTS-Oracle-predicted TREM2 hits. (A) Results of single-dose screening for TREM2 using the Fnorm readout after 5 seconds. Negative control (DMSO in assay buffer) and positive control (**T2337**) are shown as green dots, hits outside of a ten standard deviation range (shown as orange area) are displayed as blue dots. Compounds that had to be excluded after control experiments (Figure S3) are indicated as red hollow dots. Results from two independent experiments, graph created with GraphPad Prism 10. (B) Dose-response binding curves for validated TREM2 hits. Compounds were tested across an 8-point concentration series (200 µM to 0.005 µM) using Monolith X to determine binding affinities (n=2). (C) Individual dose-response binding isotherms for the seven validated TREM2 modulators. (D) Overview of HTS-Oracle workflow and performance metrics for TREM2 hit discovery.

The *K*_D_ values ranged from 6.46 µM to 180 µM (median *K*_D_ = 12.4 µM), with the most potent compound (**19B03**) demonstrating a K_D_ of 6.46 ± 1.17 µM. This represents a 3.5-fold improvement in binding affinity over our previously reported best TREM2 ligand (**T2337**, *K*_D_ = 22.4 µM)^28^, suggesting that HTS-Oracle not only enriches for hits but also prioritizes compounds with superior binding characteristics. The identification of multiple single-digit micromolar TREM2 binders (**13K04**, *K*_D_ = 8.18 µM, **3N08**, *K*_D_ = 7.38 µM) provides valuable chemical starting points for medicinal chemistry optimization and validates HTS-Oracle’s utility in early-stage TREM2 drug discovery programs. Collectively, these results demonstrate that HTS-Oracle can substantially accelerate hit identification for challenging immunomodulatory targets while simultaneously improving hit quality and reducing resource expenditure which is are key advantages in both academic and industrial drug discovery settings.

#### Case study 2: HTS-Oracle-Guided Discovery of CHI3L1 Inhibitors

A second case study to determine the performance of HTS-Oracle was performed to identify CHI3L1 hits. HTS-Oracle was trained using a library compiled from ChEMBL^31^ and published work^32–34^. The trained model screened an in-house compound library comprised of 10241 compounds and was followed by experimental validation of the predicted hits. Next, the trained HTS-Oracle model was employed to prioritize 19 potential hits which were selected for experimental validation using microscale thermophoresis (MST). In the primary MST screen conducted at 250 μM compound concentration, nine molecules initially exhibited detectable binding signals: **Z1729614993**, **Z2699161474**, **Z3393951732**, **Z2019428876**, **Z1693314177**, **Z2633381045**, **Z1823104988**, **Z2783588021**, and **Z1193339120** (Figure 4A). These compounds showed Fnorm values that deviated significantly from the non-binding compounds (black data points) and negative control (brown data point), suggesting potential CHI3L1 interaction. The positive control **G28**^35^, a known CHI3L1 inhibitor, demonstrated the expected binding behavior. However, careful analysis of the initial fluorescence intensities revealed potential confounding factors for two compounds (Figure S4).

**Figure 4.**
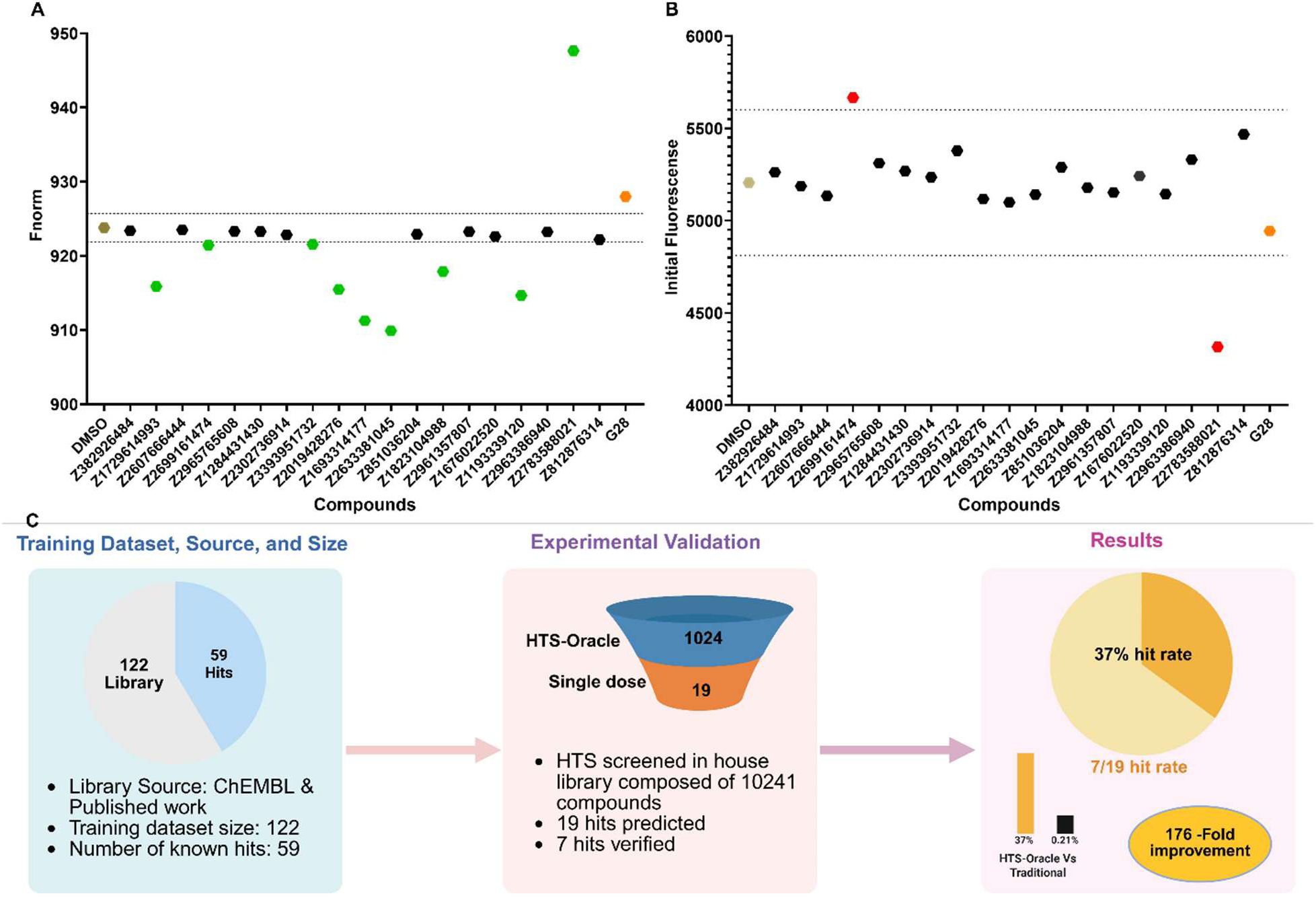
Experimental validation of HTS-Oracle-predicted CHI3L1 hits. Primary screening of compounds at 250 μM (with 3.1% DMSO). Green, brown, black, and orange indicate potential hits, negative control/reference (buffer with 3.1% DMSO), non-binders, and positive control (**G28**^35^), respectively; (B). (C) Overview of HTS-Oracle workflow and performance metrics for TREM2 hit discovery.

Among the tested compounds, **Z2699161474** exhibited substantially elevated initial fluorescence compared to the DMSO control and other test compounds, indicating intrinsic autofluorescence that could generate false-positive binding signals in the MST assay. Conversely, **Z2783588021** displayed markedly reduced initial fluorescence, characteristic of fluorescence quenching behavior. Such optical interference can compromise MST data interpretation by affecting the fluorescence-based thermophoresis readout independently of actual binding events. Follow-up orthogonal control experiments confirmed these fluorescence artifacts, necessitating exclusion of both compounds from the validated hit list.

After removing these two artifactual binders, seven compounds—**Z1729614993**, **Z3393951732**, **Z2019428876**, **Z1693314177**, **Z2633381045**, **Z1823104988**, and **Z1193339120** were confirmed as genuine CHI3L1 binders, representing a 37% experimental hit rate (7 out of 19 tested compounds). The 37% hit rate in experimental validation confirms that HTS-Oracle effectively learned the chemical features associated with CHI3L1 binding from the training dataset and successfully generalized these patterns to identify novel active compounds. More significantly, the 37% hit rate represents a 1176-fold improvement in screening efficiency compared to the traditional HTS screening with a true hit rate of 0.21%^36^ (11 hits out of 5280 tested compounds)^36^. This significant increase in hit rate reinforces the ability of the platform to substantially reduced the time, cost, and material resources required for experimental validation while maintaining high predictive accuracy.

#### Validating HTS-Oracle on Imbalanced Datasets: CD28 Inhibitors Discovery

HTS-Oracle demonstrated exceptional performance in the CHI3L1 and TREM2 hit identification campaigns, where the training datasets contained relatively balanced hit rates of 48% (59 hits out of 122 compounds) and 14% (63 hits out of 461 compounds), respectively. These balanced datasets provided the machine learning algorithm with substantial positive examples to learn the chemical features associated with target binding. However, real-world drug discovery scenarios often present a more challenging situation: identifying active compounds within large chemical libraries where the known hits represent only a small fraction of the total collection which is a condition known as severe class imbalance.

To rigorously evaluate HTS-Oracle’s capability to handle highly imbalanced datasets representative of typical screening campaigns, we carried out a third case study using a significantly more challenging training set. HTS-Oracle was trained on a library containing only 120 confirmed hits among 6,003 total compounds, representing a hit rate of just 2.0% which is substantially lower than the CHI3L1 (48%) and TREM2 (14%) training sets. This <5% hit rate reflects the reality of many high-throughput screening campaigns and represents a stringent test of the platform’s ability to extract meaningful structure-activity patterns from sparse positive training examples while avoiding false positives from the overwhelming majority of inactive compounds.

HTS-Oracle was trained on a highly imbalanced CD28-binding dataset derived from screening over 7,000 curated small molecules using orthogonal high-throughput assays^37,38^. CD28 was chosen due to its central role in T-cell activation and it’s the historical difficulty of its targeting using small molecules^39,40^. The final training set comprised 6,003 unique compounds containing only 120 confirmed hits, representing a challenging hit rate of just 2.0%. This severely imbalanced dataset, with positive examples representing less than 5% of the total library, provided a stringent test of the platform’s ability to extract meaningful structure-activity patterns from sparse positive training data while minimizing false positives from the overwhelming majority of inactive compounds.

Next, the Trained HTS-Oracle model was applied prospectively to an evaluation set of 1,152 in house compounds that were never subjected for CD28 testing. Given the imbalanced nature of the training dataset, a wider range was employed where HTS-Oracle was used to prioritize 345 molecules (1 plate) for experimental testing which is a 70% reduction in screening burden. Compounds were screened in 384-well plates at a concentration of 200 µM using a PBS-based assay buffer. Hit selection was based on ΔF_norm_ values derived from TRIC measurements and compounds exceeding three times the standard deviation of the negative control were considered to be hits. Compounds exhibiting autofluorescence, quenching, or aggregation artifacts were excluded.

Based on these selection criteria, 29 primary hits were identified from the computationally focused library of 345 compounds, yielding a remarkably high hit rate of 8.41% (Figure 5). When benchmarked against the original unfiltered library of 1,120 compounds, our trained model achieved a dramatic 70% reduction in the experimental screening effort while simultaneously delivering an 8-fold enhancement in hit rate compared to previous CD28-targeted HTS campaigns (Figure 5). The 8.41% hit rate achieved through this focused screening represents a marked departure from the typical 1%^8,41^ hit rates commonly observed in HTS campaigns. Together, these results highlight the ability of HTS-Oracle in effectively handling severely imbalanced datasets typical of real-world drug discovery while maintaining high predictive accuracy and substantially improving screening efficiency.

**Figure 5.**
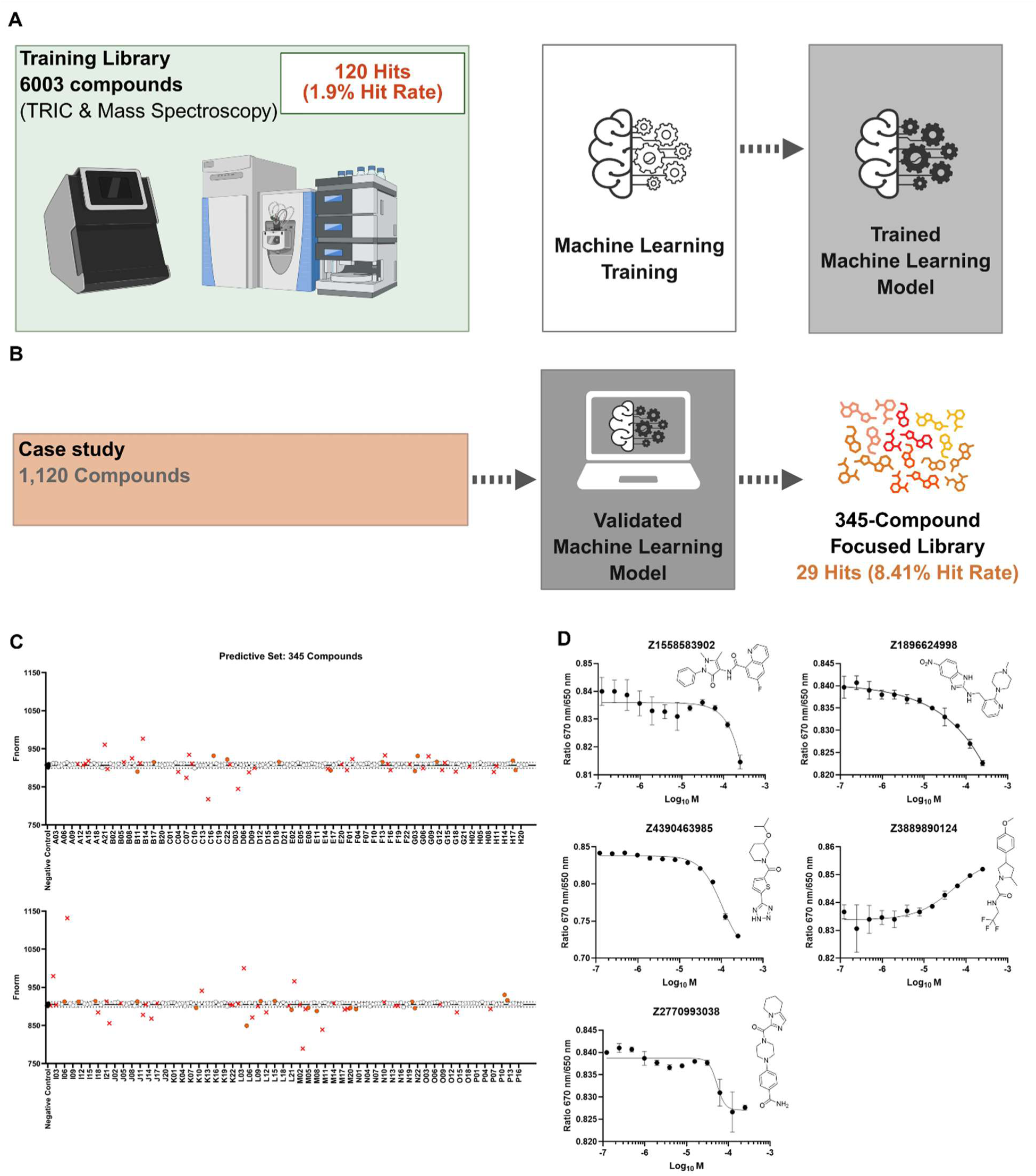
HTS-Oracle workflow and performance for CD28 hit discovery from a highly imbalanced dataset. (A-B) Schematic overview of the training, validation, and application of the ML model. (C) Primary screening results of the 345 HTS-Oracle-predicted compounds using TRIC assay. (D) Binding affinity measurements of test compounds to CD28 protein using spectral shift assay.

Among the 29 hits identified in the TRIC-based screening, 14 compounds were commercially available and subsequently selected for binding affinity measurements using spectral shift detection. Of these, five compounds showed dose-dependent binding affinity toward CD28. Notably, three compounds exhibited quantifiable binding interactions with the CD28 protein (Figure 5). Compound **Z2770993038** demonstrated the highest affinity, with a dissociation constant (*K*_D_) of 43.35 ± 16.44 μM, followed closely by **Z4390463985** (*K*_D_ = 45.76 ± 12.06 μM) and **Z3889890124** (*K*_D_ = 49.53 ± 15.3 μM). All three compounds produced characteristic dose-dependent binding curves with saturation behavior, confirming specific and saturable interactions with CD28 (Figure 5). Conversely, **Z1558583902** and **Z1896624998** showed limited binding responses within the tested concentration range, suggesting potential binding interactions that may occur at higher concentrations.

Given the well-established role of CD28 in T cell co-stimulation and the significant therapeutic potential of targeting the CD28–B7.1 interaction for modulating immune responses in autoimmune diseases, transplant rejection, and cancer immunotherapy, we evaluated the functional impact of the top two hits on CD28–B7.1 binding using an ELISA-based assay^42–44^. CD28 delivers essential co-stimulatory signals that enhance T cell proliferation and drive the secretion of pro-inflammatory cytokines such as IL-2, IFN-γ, and TNF-α^45–47^. To investigate the ability of small molecules to block CD28–B7.1 binding, we performed a competitive ELISA using immobilized CD28 and biotinylated B7.1 ^48^. Two compounds, **Z2770993038** and **Z4390463985**, were evaluated across a concentration range. Both compounds demonstrated dose-dependent inhibition of CD28-B7.1 binding, with IC₅₀ values of 32.87 ± 5.74 µM and 56.29 ± 11.24 µM, respectively (Figure 6). These functional results correlate with their binding affinity obtained from the MST assay, confirming them as CD28 binders. Overall, these findings confirm that both compounds directly target CD28 and can disrupt its interaction with B7.1, supporting their potential as early-stage small molecule inhibitors for further lead optimization and therapeutic development.

**Figure 6.**
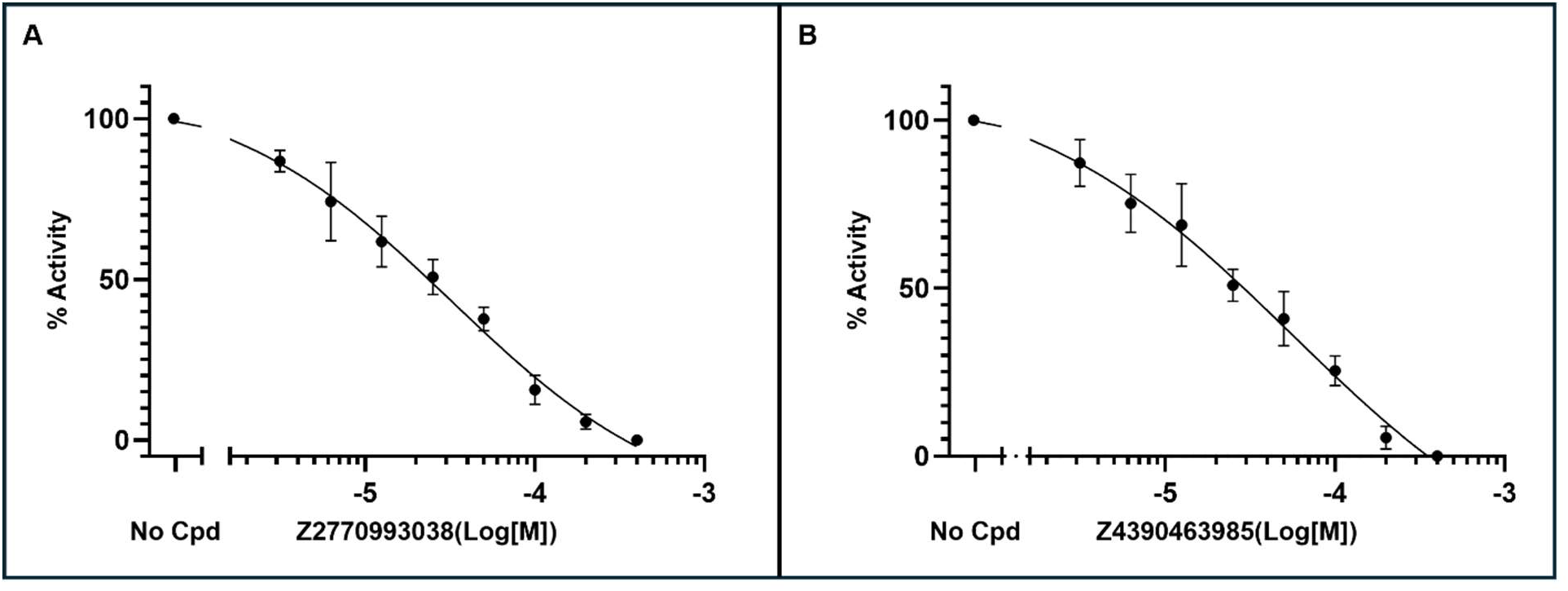
Dose-dependent inhibition of CD28–B7.1 interaction by compounds. The ability of compounds, **Z2770993038** and **Z4390463985** to inhibit CD28–B7.1 binding was assessed using a competitive ELISA assay. IC₅₀ was determined using GraphPad Prism with a four-parameter variable slope nonlinear regression model. Data represent mean ± standard deviation from 3 independent repeats.

## Methods

### Model Development

A hybrid deep learning framework was developed to identify active compounds from HTS libraries by combining chemical structure-based representations with ensemble classification. Compounds were obtained from library.csv (containing SMILES strings and compound IDs) and positives.csv (containing confirmed actives). Each compound was assigned a binary activity label, where y=1, y=1 if its ID appeared in the positives set and y=0, otherwise.

Each SMILES string was featurized using three complementary molecular representations: Morgan circular fingerprints^49^ were computed with radius 2 and 2,048 bits, encoding neighborhood substructures. In parallel, 167-bit MACCS keys^50^ were generated to capture common substructural motifs. Additionally, a set of 15 physicochemical descriptors was computed using RDKit^16^, including molecular weight, Log*P*, topological polar surface area, hydrogen bond donors and acceptors, rotatable bonds, ring count, and QED. These vectors were concatenated to form a 2,230-dimensional feature vector per compound.

To reduce dimensionality, we applied three feature selection methods independently: LASSO regression^51^, principal component analysis (PCA)^52^, and mutual information ranking^53^. LASSO performs feature selection by minimizing the L1-penalized loss:

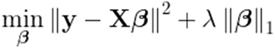

For PCA, features were projected onto principal components, and the top 200 were retained. Mutual information selection ranked features by their dependency with the activity labels. Subsequently, a dual-branch neural network was trained using five-fold stratified cross-validation. One branch used ChemBERTa^15^, a transformer-based model pretrained on SMILES, to encode each molecule’s SMILES into a dense vector representation. The second branch processed the selected RDKit features through a multilayer perceptron. Outputs from both branches were concatenated and passed through fully connected layers to produce a final prediction score. Training used the AdamW optimizer and binary cross-entropy loss with class weights to address data imbalance:

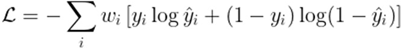

where *ŷ_i_* is the predicted probability and *w_i_* is the class weight.

Model predictions were averaged across folds and feature selection strategies. For each compound *i*, the final ensemble prediction was computed as the mean of its valid model outputs:

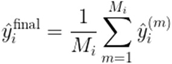

Model performance was assessed using the area under the receiver operating characteristic curve (ROC AUC), average precision (AP), precision, recall, and F1-score among other features. The script employed to evaluate the trained ML model “evaluation.py” is available in the GitHub repository. Final prediction scores were exported for downstream prioritization.

### Web Application Development

A Streamlit application was developed to implement the trained ensemble model for predicting molecular activity in new datasets, offering a user-friendly interface for researchers. The application allows users to upload molecular libraries in CSV format and receive predictions in a streamlined workflow. To ensure compatibility with diverse data sources encountered in chemical and pharmaceutical research, the application automatically identifies SMILES columns through pattern matching against standard naming conventions. For each molecular input, the application constructs a comprehensive descriptor-based representation. This feature generation step includes Morgan fingerprints, MACCS keys, and physicochemical descriptors. Morgan fingerprints are calculated from the SMILES representation using the equation:

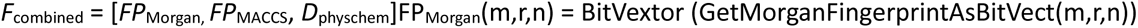

where *m* is the molecular structure, *r* = 2 the radius, and *n* = 2048 n=2048 is the fingerprint bit length. The final feature vector contains molecular properties such as molecular weight, Log*P*, rotatable bonds, hydrogen bond acceptors/donors, topological polar surface area, aromatic rings, heteroatoms, sp³ carbon fraction, and QED score.

The application uses an ensemble-based classification framework built on multiple ML models trained with different feature selection strategies, namely LASSO, PCA, and mutual information. For each method 𝓀, predictions are averaged across models using:

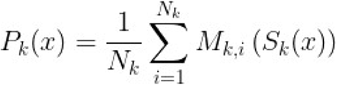

where *ℳ*_𝓀,*i*_ is the *i^th^* model for method 𝓀, and 𝒮_𝓀_(𝓍) is the feature-selected transformation of the input vector 𝓍. The final ensemble prediction is obtained by averaging over all methods:

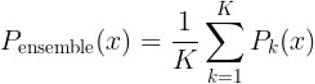

with K=3 in. Predictions are then converted into binary activity labels using a classification threshold θ (default = 0.5), defined as:

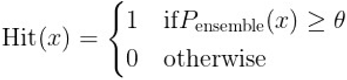

In addition to classification, the model also outputs prediction confidence, calculated as 2⋅∣P_ensemble_ (x) −0.5 ∣, and ensures that all scores are clipped to the range [0,1] to maintain probabilistic interpretability.

To ensure reliability, SMILES validation is performed using RDKit parsing before feature extraction^54^. Descriptors are also sanitized by replacing NaN or infinite values with zeroes. The application supports efficient batch processing with real-time progress updates and robust error tracking. Finally, prediction scores are visualized using interactive, color-coded tables where color intensity scales linearly with predicted activity. This visualization aids users in rapidly identifying top candidates for downstream experimental validation.

### TREM2 case study methodology

#### Screened library

The screened library comprised an in-house collection of 12,712 compounds which were purchased from Enamine.

#### Single-dose screening for TREM2

The single-dose screening was conducted as described previously^28^ using recombinant human His-tagged TREM2 (SinoBiological, #11084-H08H) labeled with RED-tris-NTA 2^nd^ generation dye (NanoTemper Technologies, #MO-L018). His-labeled TREM2 (10 nM) was then incubated with the selected compounds (100 µM) in assay buffer (PBS, pH 7.4, 0.05% Tween20, 2% final DMSO) for 15 minutes at room temperature. A negative control (n = 8 per replicate) consisting of assay buffer was run alongside the samples as well as a positive control (**T2337**^28^, 100 µM, n = 8 per replicate). The plates were measured on a Dianthus NT.23Pico (NanoTemper Technologies) using TRIC (5s laser time). The results from two independent experiments were analyzed based on the normalized fluorescence (F_norm_) for all wells after 5 s laser time. Compounds were considered potential hits if their average F_norm_ was outside of a ten standard deviation range from the negative control. To exclude assay interfering compounds, we incubated all compounds with assay buffer and RED-Tris-NTA 2^nd^ generation dye instead of labeled protein and measured their initial fluorescence and TRIC signal with Dianthus (Figure S3, Table S1). Compounds that were flagged by the software (DI.Analysis) as aggregators or that displayed a significant difference (≥20%) in initial fluorescence from the average reference (n = 4) were excluded. The data were plotted in GraphPad Prism 10 for visualization.

#### TREM2 binding affinity

The experiments were performed on a Monolith X (NanoTemper Technologies) using Monolith Premium Capillaries (NanoTemper Technologies, #MO-K025). His-tagged TREM2 (SinoBiological, #11084-H08H) was labeled with RED-tris-NTA 2^nd^ generation dye (NanoTemper Technologies, #MO-L018) according to the manufacturer’s instructions and the protein concentration was adjusted to 40 nM. The ligand solutions were prepared from the 10 mM stock on the library plates as 2-fold stocks at 400 µM in 2-fold assay buffer (PBS, pH 7.4, 0.05% Tween20, 10% DMSO) and a 16-point dilutions series (1:1) was prepared. Then, all samples were incubated with a 2-fold protein stock (40 nM) in PCR tubes for 15 min at room temperature (final concentrations: 200–0.005 µM ligand, 20 nM His-TREM2, 5% DMSO). Capillaries were dipped into each tube, loaded onto the chip, and the measurement was started at 100% excitation with medium MST power.

Thermophoresis was measured for 10 seconds with an initial five second delay. All samples were run in two independent experiments. The raw TRIC data were analyzed using the instrument’s software (MO.Control 2) to create merge sets of all replicates per compound. Capillaries that were flagged by the software as initial fluorescence outliers or for aggregation were excluded from the analysis. For better comparability between different compounds, data were normalized to fraction bound. The data were plotted in GraphPad Prism 10 for visualization.

### CHI3L1 case study methodology

Protein–ligand interactions were evaluated by microscale thermophoresis (MST) using our previously described protocol. CHI3L1-His was fluorescently labeled with RED-tris-NTA dye (NanoTemper Technologies) following the manufacturer’s instructions. The labeled protein was incubated with test compounds under optimized assay conditions and measurements were performed on the Dianthus NT.23 Pico system.

To account for potential assay artifacts arising from compound-dependent autofluorescence or fluorescence quenching, dedicated control experiments were conducted. For autofluorescence, fluorescence of the labeled protein in 3.1% DMSO was compared with that of buffer containing 250 μM compound in 3.1% DMSO. For quenching assessment, fluorescence from 20 nM dye in buffer containing 3.1% DMSO was compared with that of 20 nM dye incubated with each compound under same solvent conditions. Compounds exhibiting fluorescence values outside the mean ± 3 SD of the respective controls were excluded from further analysis.

### CD28 case study methodology

#### Single-dose screening Using TRIC Technology

Evaluation of the HTS-Oracle prioritized compounds for CD28 binding was conducted using the Dianthus NT.23 Pico system (NanoTemper Technologies)^37^. Selection criteria were applied as previously outlined^55^. Briefly, compounds that produced a ΔF_norm_ greater than three times the standard deviation of the negative control were initially considered potential CD28 binders. After excluding compounds that exhibited autofluorescence, quenching, aggregation, or scan anomalies, the remaining compounds were classified as primary hits. An additional control experiment was conducted for each selected small molecule to ensure no interference with the RED-tris-NTA dye alone - either by quenching its signal, increasing its fluorescence intensity, or altering the TRIC trace. Compounds that exhibited any interaction with the dye alone were excluded from further analysis. The final set of small molecules was then selected for binding affinity measurements and further downstream validation experiments.

#### 4.5.2. MST

Binding experiments were conducted in assay buffer containing 154 mM NaCl, 5.6 mM Na₂HPO₄, 1.05 mM KH₂PO₄, pH 7.4, and 0.005% Tween-20. For binding measurements, labeled CD28 protein was used at a final concentration of 40 nM, and RED-tris-NTA 2nd Generation was added at 20 nM. Test compounds were prepared in DMSO and added to achieve a final DMSO concentration of 1% (v/v). The mixture was incubated for 10 minutes at room temperature prior to measurements.

The Monolith X instrument was configured with optimized parameters for spectral shift detection. All measurements were performed in triplicate using premium capillaries. Initial data processing was performed using the MO. Affinity Analysis software (NanoTemper Technologies), and subsequent data analysis and curve fitting were conducted using GraphPad Prism (GraphPad Software, San Diego, CA, USA).

#### 4.5.3. ELISA

A competitive ELISA was carried out using a commercially available kit from BPS Bioscience (CD28:B7.1[Biotinylated] Inhibitor Screening Assay Kit) to evaluate the ability of test compounds to inhibit the binding of biotinylated B7.1 to immobilized CD28^48^. The experiment was performed following the protocol provided with the kit, and a brief description of the procedure is provided below.

Ninety-six–well plates were coated with CD28 protein at a concentration of 2 µg/mL in PBS (50 µL per well) and incubated overnight at 4°C. The following day, wells were washed with 1× Immuno Buffer and blocked with Blocking Buffer for 1 hour at room temperature to minimize nonspecific binding. Test compounds were added at various concentrations together with biotinylated B7.1 (5 ng/µL), and the mixture was incubated for 1 hour to facilitate competitive binding. Uncoated wells served as negative controls for background signal, while wells containing inhibitor buffer in place of compound served as assay controls. After the binding step, wells were washed thoroughly, followed by the addition of Streptavidin-HRP (diluted 1:1000 in Blocking Buffer) and incubation for 1 hour at room temperature. After a final wash, a chemiluminescent substrate was added, and the luminescence signal was measured immediately using a plate reader in luminescence detection mode.

### Practical Guidelines and considerations for the Use of HTS-Oracle

HTS-Oracle was developed to enhance compound prioritization in HTS campaigns by improving predictive performance and reducing experimental burden. While the model demonstrated strong performance in both retrospective and prospective validations, its successful implementation depends on careful consideration of several practical and methodological factors.

HTS-Oracle is not a universally applicable model; the model must be retrained for each new biological target using a representative and curated training dataset. Its architecture is target-agnostic, but the predictive utility depends critically on the underlying data. Nonetheless, the pipeline is computationally efficient. For example, inference across a library of 6,000 compounds can be completed in under ten minutes on a workstation equipped with an NVIDIA RTX 4070 Ti GPU and a 32-core CPU, allowing rapid deployment once model training and validation are complete.

To minimize false positives, which can be costly in follow-up experimental assays, HTS-Oracle employs a conservative prediction strategy that prioritizes high precision. This design choice reduces the risk of pursuing inactive compounds but may also increase false negatives, particularly for compounds with borderline activity scores. This trade-off is especially acceptable in low hit-rate settings, such as the CD28 campaign described here (∼1-2%), where experimental validation is resource-intensive and the cost of false positives is high. In contrast, screens with higher baseline hit rates (such as TREM2 and CHI3L1) benefited from adjusted prediction thresholds to maximize recall.

To facilitate community adoption, HTS-Oracle is released as open-source, including both application and training code, along with a pre-trained model available through our GitHub repository. Future work will expand its application to additional target classes and assay types, refine model calibration under varying hit-rate conditions, and explore its integration with orthogonal data modalities such as transcriptomic or structural information. Ongoing updates will aim to support broader reproducibility, adaptability, and sustained model performance across diverse screening contexts.

## Conclusions

In this study, we introduce HTS-Oracle, a retrainable, open-source deep learning–based platform for hit prioritization that integrates multi-modal molecular representations in a unified ensemble framework. HTS-Oracle decreased the number of compounds requiring laboratory testing by more than 99% for TREM2 and CHI3L1 and by 70% for CD28, highlighting its capacity to prioritize hits with minimal wet-lab effort. Correspondingly, enrichment factors ranged from 8.4× to 176× relative to traditional HTS, underscoring robust target-specific performance even under challenging data conditions such as the highly imbalanced CD28 training set. Beyond, proving the success of the platform, we report several novel and potent leads for three therapeutic targets which have been considered “undruggable” and should be considered for further optimization. HTS-Oracle addresses two persistent bottlenecks in drug discovery: the limited success of traditional HTS campaigns and the difficulty of identifying chemically diverse yet potent starting points. By leveraging both contextual and structural features of molecules, the model navigates broad chemical space while maintaining selectivity and interpretability—characteristics essential for translation into medicinal chemistry workflows.

Looking forward, we anticipate that HTS-Oracle will serve as a generalizable blueprint for AI-enhanced virtual screening across a wide range of therapeutic targets. Its open-source availability, modular architecture, and demonstrated experimental validation provide a strong foundation for adoption across academic and industry settings. Future extensions may include integration with protein structural data, bioactivity profiles, transcriptomic signatures, or multitask learning frameworks to further enhance predictive performance and biological relevance. In conclusion, HTS-Oracle represents a scalable, experimentally grounded, and practically deployable platform that bridges the gap between large-scale virtual screening and focused, resource-efficient hit validation. Its successful application to CHI3L1, TREM2 and CD28 highlights the potential of machine learning to redefine early-phase drug discovery—making previously inaccessible targets newly tractable.

## Acknowledgments

This work was supported by the National Institute of Diabetes and Digestive and Kidney Diseases (NIDDK) under grant number R01DK137299.

## Data Availability Statement

All datasets used for model development and validation are publicly available in our GitHub repository at https://github.com/HTS-Oracle/. Supplementary figures and tables are provided in the Supplementary Information.

## Code availability

❖ HTS-Oracle is freely available, including environment setup instructions, source code, system requirements, trained models, and a Streamlit web application. All resources can be accessed via GitHub at https://github.com/HTS-Oracle/HTS-Oracle-TREM2.
❖ Pre-trained models for TREM2, CHI3L1, and CD28 targets are available for immediate use without retraining at https://osf.io/3nj8t.

