## Supporting Information for "HTS-Oracle: Experimentally validated AI-enabled prioritization for generalizable small molecule hit discovery"

**Appendix**

| Figures S1-4 | Pages 2-3 |
| --- | --- |
| Table S1 | Page 4 |

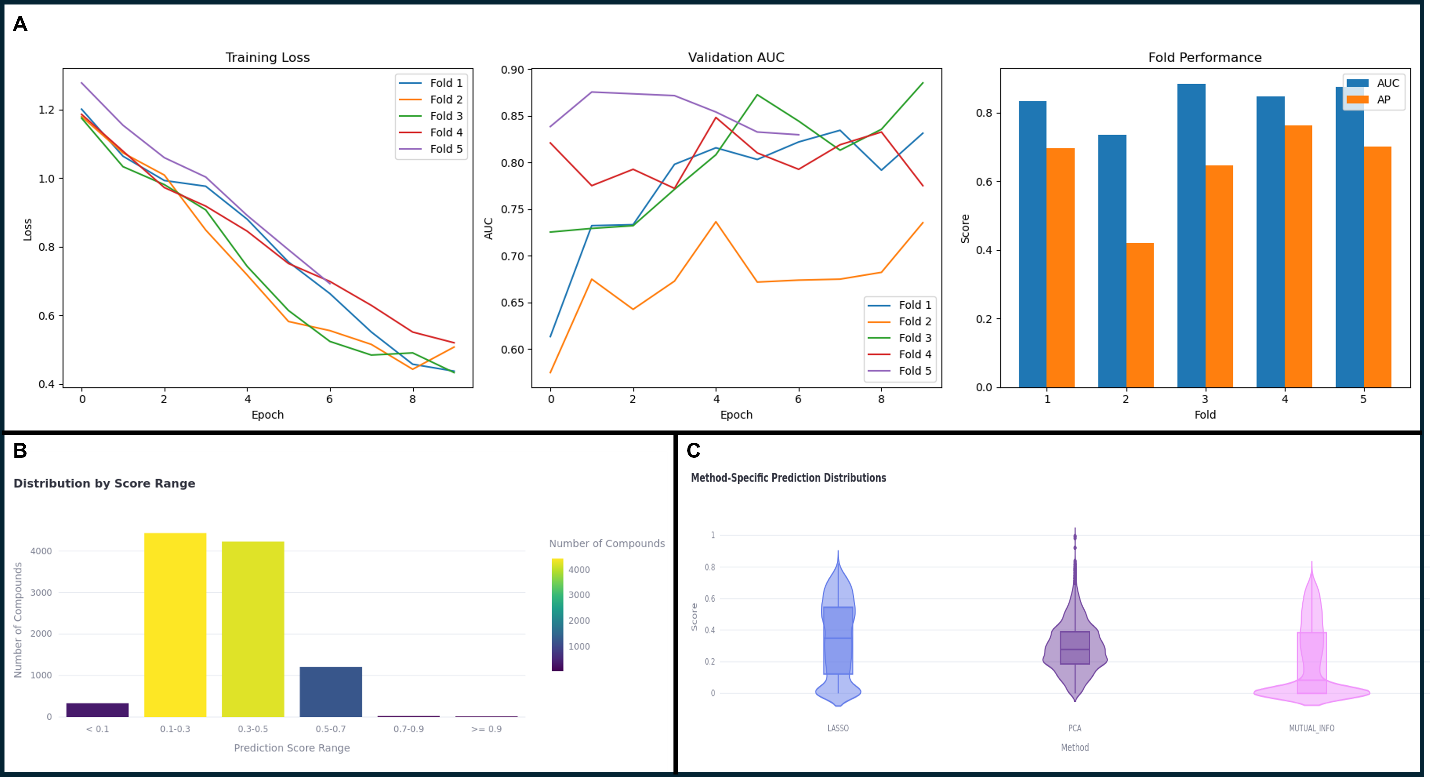

**Figure S1. Cross-validation performance and prediction distribution analysis of HTS for TREM2 hit prediction.** (A) Model training and validation performance across 5-fold cross-validation*. Left panel*: Training loss curves showing convergence across all folds over 10 epochs. *Middle panel*: Validation AUC trajectories demonstrating model stability and performance during training. *Right panel*: Final performance metrics (AUC and Average Precision, AP) for each fold, with mean AUC = 0.83 ± 0.06 and mean AP = 0.61 ± 0.15. (B) Distribution of compounds by prediction score range. (C) Method-specific prediction score distributions comparing three approaches: LASSO (blue), PCA (purple), and MUTUAL_INFO (pink).

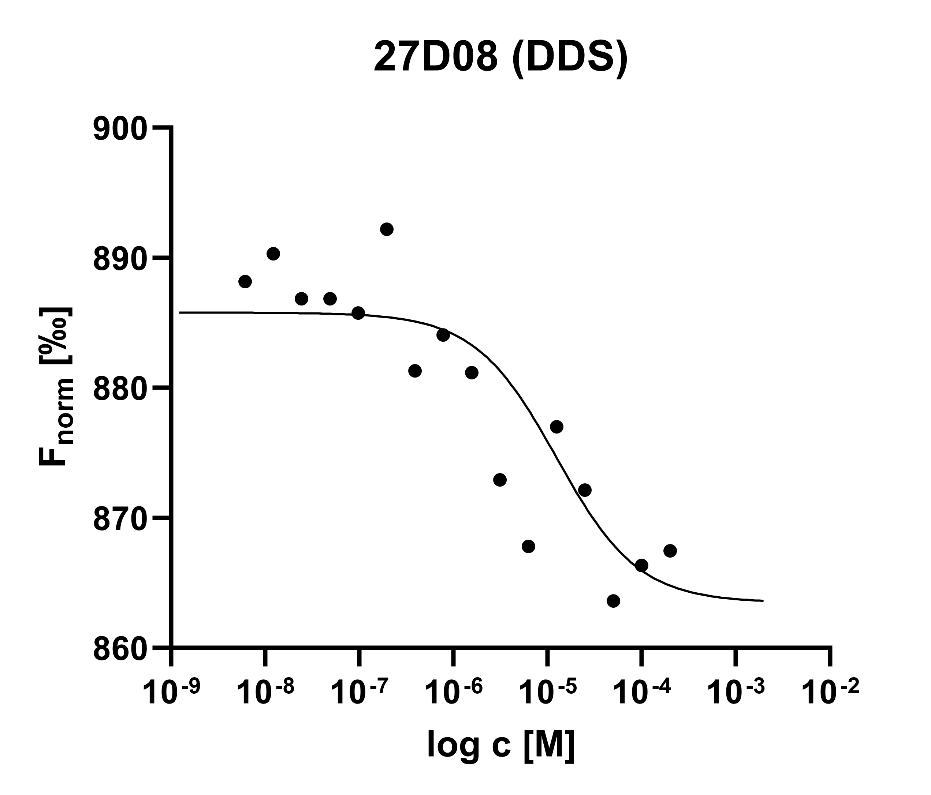

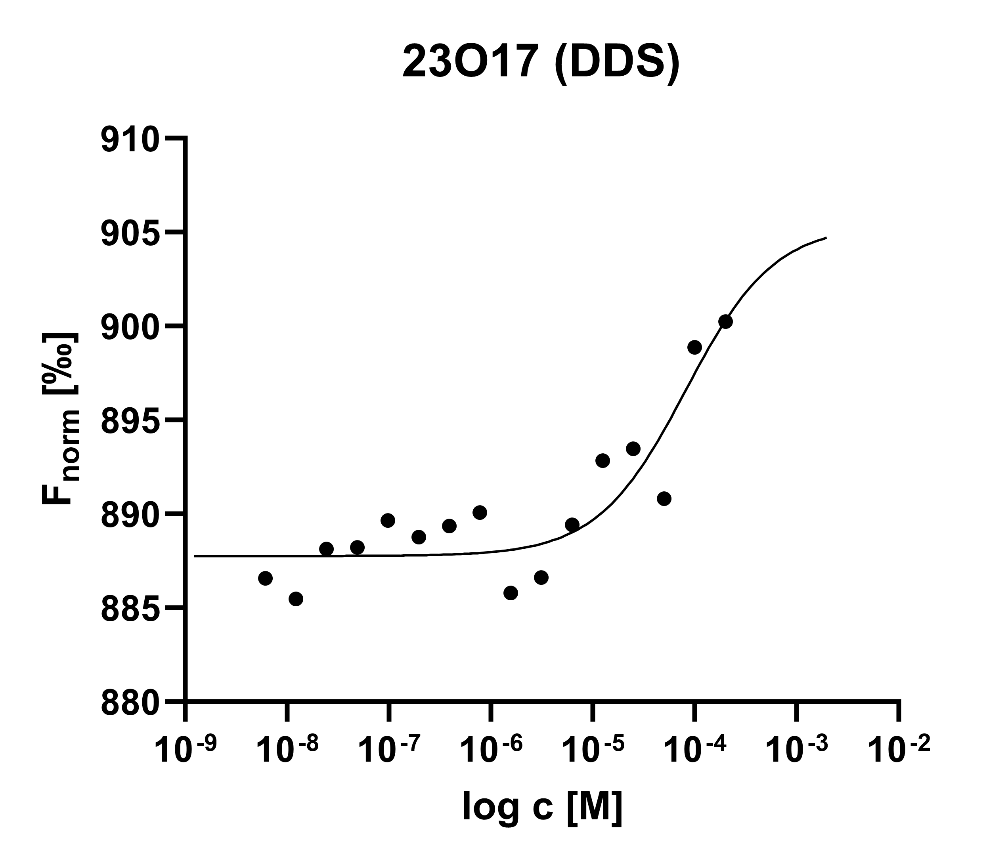

**Figure S2.** F_norm_ vs log c plots for hits from HTS-Oracle measured on Monolith X. Graphs created with GraphPad Prism 10 as merge sets from n=1 due to limited availability on stock plate).

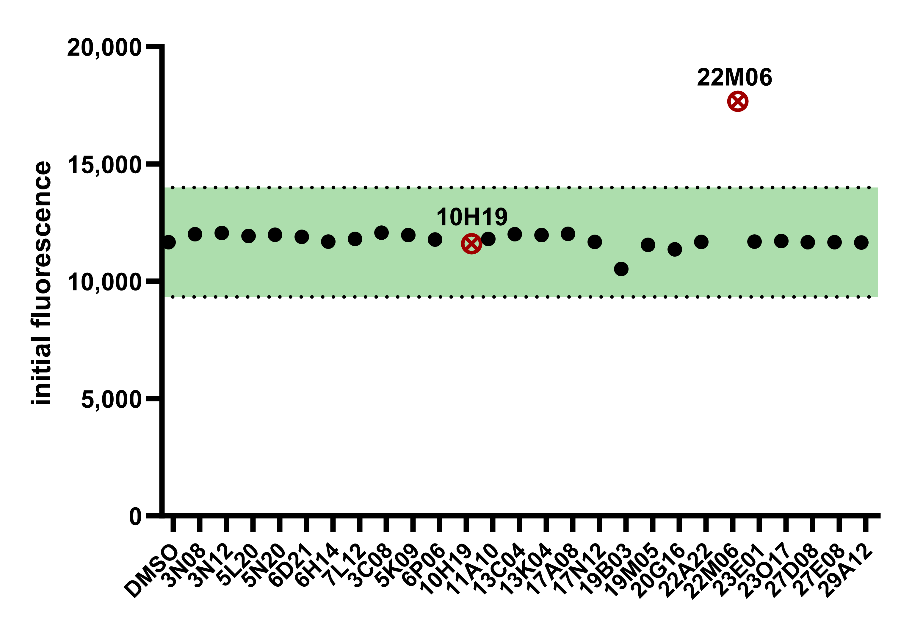

**Figure S3.** Results of control experiments. Compounds that were outside of a 20% range (22M06) from the initial fluorescence of the average reference (n = 4) or that showed aggregation (10H19) were excluded as hits.

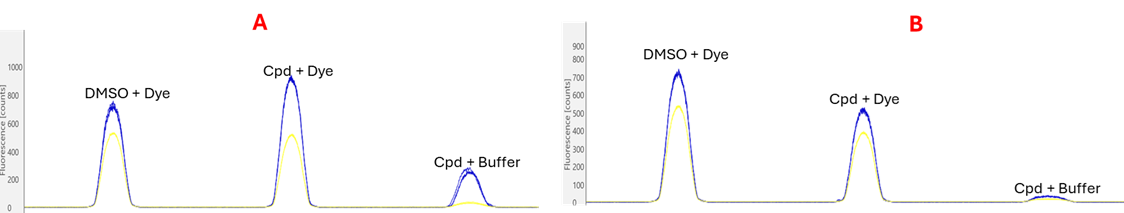

**Figure S4.** Assessment of potential artifacts from autofluorescence and quenching in MST assays by compounds **Z2699161474 (A)** and Z2783588021 **(B)**. Quenching was assessed by comparing fluorescence of 20 nM dye in buffer with DMSO to that of 20 nM dye incubated with each compound. Autofluorescence of test compounds was evaluated by comparing the fluorescence of labeled protein in 2.5% DMSO with buffer containing 250 μM compound in 2.5% DMSO.

**Table S1. TREM2 screening results**

| **Catalog ID** | **screening ID** | **Plate_ID** | **Well** | **Library** | **SMILES** | **Prediction_Score** | **Confidence** | **comment** |
| --- | --- | --- | --- | --- | --- | --- | --- | --- |
| Z4252262326 | 6P06 | 1953578-Y10-06 | P06 | DDS | O=C(NC1=NNC=2C=CN=CC12)C=3C=CC=C4NC(=O)COC34 | 0.94301033 | 0.88602066 | non-binder |
| **Z2212656233** | **27D08** | 1953578-Y10-27 | D08 | DDS | O=C(NCC=1C=CC(=CC1)C(=O)N2CCOCC2)N3CCC4=C(C3)N=NN4C=5C=CC=CC5 | 0.938389599 | 0.876779199 | hit |
| Z2186482426 | 17N12 | 1953578-Y10-17 | N12 | DDS | O=C(NCCCC1=NC=CS1)N2CCC(CC2)C3=NC(=NO3)C=4C=CC=CN4 | 0.93819344 | 0.876386881 | non-binder |
| **Z1421216186** | **20G16** | 1953578-Y10-20 | G16 | DDS | CCOC=1C(Cl)=CC(=CC1OC)C(=O)NC(C)C2=NOC(=N2)C=3C=CC=CC3 | 0.937030494 | 0.874060988 | hit |
| Z3268551692 | 22A22 | 1953578-Y10-22 | A22 | DDS | CCN1C=C(NC(=O)NCCNC=2C=CC=CC2)C(=N1)C(=O)N | 0.929047823 | 0.858095646 | non-binder |
| Z3464156438 | 22M06 | 1953578-Y10-22 | M06 | DDS | OC1CCCC=2NC(=O)C(=CC12)C(=O)NC=3C=CC(O)=CC3Cl | 0.928589761 | 0.857179523 | assay interference |
| **Z3417380680** | **23O17** | 1953578-Y10-23 | O17 | DDS | CN1C=CC=2C=CN=C(C(=O)NCC=3C=CC=4CN(C5CCC(=O)NC5=O)C(=O)C4C3)C12 | 0.926834106 | 0.853668213 | hit |
| **Z2065713205** | **13C04** | 1953578-Y10-13 | C04 | DDS | CC1=CC=2C=C(CNC(=O)C=3C=CC=C(C3)[N+](=O)[O-])C=CC2N1 | 0.922432601 | 0.844865203 | hit |
| Z1928169577 | 27E08 | 1953578-Y10-27 | E08 | DDS | CN1C(=CC=2C(=O)N(C)C(=O)N(C)C12)C(=O)NCCC3=NC(=O)ON3 | 0.922411621 | 0.844823241 | non-binder |
| Z4252257848 | 29A12 | 1953578-Y10-29 | A12 | DDS | O=C(NCC=1C=CC=2NC(=O)NC2C1)C3CC(=O)NC=4N=CC=CC34 | 0.921911716 | 0.843823433 | non-binder |
| Z813469362 | 23E01 | 1953578-Y10-23 | E01 | DDS | CC=1C=C(C=CC1NC(=O)N2CCC=3NC=4C(Cl)=CC(Cl)=CC4C3C2)C(=O)N | 0.916840672 | 0.833681345 | non-binder |
| Z1245744325 | 10H19 | 1953578-Y10-10 | H19 | DDS | C=1OC(=NC1C2=NC(=NO2)C=3C=CC=CN3)C=4C=CC=CC4 | 0.9149459 | 0.829891801 | aggregator |
| **Z4866922164** | **3C08** | 1953578-Y10-03 | C08 | DDS | O=C(CN1C(=O)NC(C2CC2)(C1=O)C3=CC=CS3)NC(CC=4C=CC=CC4)C(=O)NC5CC5 | 0.914171159 | 0.828342319 | hit |
| **Z2444355017** | **19B03** | 1953578-Y10-19 | B03 | DDS | C(OC=1C=CC(CC2=NOC(=N2)C3CCC=4N=CN=CC4C3)=CC1)C=5C=CC=NC5 | 0.909697533 | 0.819395065 | hit |
| **Z68482620** | **17A08** | 1953578-Y10-17 | A08 | DDS | COC=1C=CC=CC1NS(=O)(=O)C=2C=C(C=CC2C)C(=O)NCC(O)C=3C=CC=CC3 | 0.907739103 | 0.815478206 | hit |
| **Z6619029074** | **13K04** | 1953578-Y10-13 | K04 | DDS | NC(=O)NC(=O)CCNC(=O)C=1N=C(N)C=C2C=CC=CC12 | 0.843021274 | 0.686042547 | hit |
| **Z3417378782** | **19M05** | 1953578-Y10-19 | M05 | DDS | NC=1N=C(N=CC1C(=O)NCC=2C=CC=3CN(C4CCC(=O)NC4=O)C(=O)C3C2)C5CC5 | 0.826937497 | 0.653874993 | hit |
| Z4136868486 | 5K09 | 1953578-Y10-05 | K09 | DDS | CC=1N=C(F)C=CC1NC(=O)NC2CCCC=3SC(=NC23)C=4C=CC(F)=CC4 | 0.8 | 0.6 | non-binder |
| Z336089236 | 6H14 | 1953589-Y10-06 | H14 | glycomimetics | CC(=O)NC1C(O)C(O)C(CO)OC1OC=2C=CC=C3C=CC=NC23 | 0.920156479 | 0.840312958 | non-binder |
| **Z1784077269** | **3N08** | 1953589-Y10-03 | N08 | glycomimetics | CC(=O)C=1C=CC(O[C@@H]2O[C@H](CO)[C@H](O)[C@H](O)[C@H]2O)=CC1 | 0.908093989 | 0.816187978 | hit |
| Z1346385512 | 5L20 | 1953589-Y10-05 | L20 | glycomimetics | CC1(O)CC2CCC(C1)N2 | 0.707787395 | 0.415574789 | non-binder |
| Z1383883638 | 7L12 | 1953589-Y10-07 | L12 | glycomimetics | NC(=O)C1CCCC1NC=2N=C(CN3CCOCC3)N=C4SC=C(C=5C=CC=CC5)C24 | 0.632809015 | 0.265618031 | non-binder |
| **Z56777074** | **5N20** | 1953589-Y10-05 | N20 | glycomimetics | CC(=O)OCC1OC(C(OC(=O)C)C1OC(=O)C)N2C(=NC=3C(N)=NC=NC23)S(=O)(=O)C | 0.619722381 | 0.239444761 | hit |
| Z494682510 | 6D21 | 1953589-Y10-06 | D21 | glycomimetics | CS(=O)(=O)N1CCC(CC1)NC(=O)NC2CCCCNC2=O | 0.618251812 | 0.236503624 | non-binder |
| Z1568353210 | 3N12 | 1953589-Y10-03 | N12 | glycomimetics | NC(=O)NC1CCN(CC2=CC(=CS2)C=3C=CC=CC3)CC1 | 0.615480065 | 0.230960131 | non-binder |
